# Evidence of sinks and sources in the PLC activated PIP_2_ cycle

**DOI:** 10.1101/183509

**Authors:** Rohit Suratekar, Padinjat Raghu, Sandeep Krishna

**Affiliations:** Simons Centre for the Study of Living Machines, National Centre for Biological Sciences, TIFR, Bengaluru 560065, India; National Centre for Biological Sciences, TIFR, Bengaluru 560065, India

## Abstract

In many eukaryotic signalling cascades, receptor mediated phospholipase C (PLC) activity triggers phosphatidylinositol 4,5 bisphosphate (PIP_2_) hydrolysis leading to information transfer in cells. Coupled with PLC activation is a sequence of reactions spread across multiple compartments by which PIP_2_ is resynthesized, a process essential to maintain PIP_2_ levels and support sustained PLC signalling. The biochemical strategies to co-ordinate these reactions and support PIP_2_ levels have remained poorly understood. In particular, the question of whether the PIP_2_ cycle is a closed cycle with no net addition or loss of metabolites has not been addressed. Using mathematical modelling approaches, we find that a closed PIP_2_ cycle cannot explain experimentally observed changes in the metabolic intermediates when changing enzyme activities in the PIP_2_ cycle. Thus, we propose that the PIP_2_ cycle likely includes at least one metabolic source and one sink whose net activity results in the experimentally observed regulation of this key signalling pathway.

## Introduction

The detection of environmental changes and responding to these is fundamental to the biology of cells. This process is mediated by receptors that are coupled by cellular response effectors by signal transduction pathways. Many divergent signalling systems exist and these are often conserved during evolution. The simplest of these are the so called two component systems in bacteria where cell surface membrane receptors are coupled to response pathways by the activity of a cytoplasmic kinase. In eukaryotic cells, elaborate multiple component systems are found where, in addition to protein kinases, information transfer properties are underpinned by the generation of small diffusible chemical messengers. Examples of these include the generation of cyclic nucleotides and Ca^2+^, NO and CO. Such molecules diffuse through the aqueous cellular cytoplasm, bind to effectors and hence mediate information transfer. The activity of such messengers is terminated by mechanisms such as degrading enzymes, transporters, etc, that remove them from the activity field of their effector proteins.

In addition to water-soluble small molecules, eukaryotic cells also employ lipid molecules as second messengers. These lipids partition into cellular membranes and in contrast to structural lipids are often found at very low levels. Their levels are determined by signal regulated activity of enzymes that either synthesize or degrade them. Phosphoinositides, phosphorylated derivatives of phosphatidylinositol (PI), are a well-known example of lipid second messengers. The most abundant of these, phosphatidylinositol 4,5 bisphosphate [denoted PIP_2_ in this manuscript] is a key chemical intermediate in cellular signal transduction. Many cell surface receptors transduce the information of a ligand bound state into downstream signals by the activation of phospholipase C (PLC) enzymes at the plasma membrane; PLC catalyses the hydrolysis of PIP_2_ to generate the membrane bound lipid diacylglycerol (DAG) and water soluble inositol 1,4,5 trisphosphate (IP_3_) both of which have well known target proteins through which they transfer information [3]. Following these events, DAG and IP_3_ are both subject to multi-step metabolic conversion leading to the resynthesis of PIP_2_ (Fig 1 (A))

**Figure 1.**
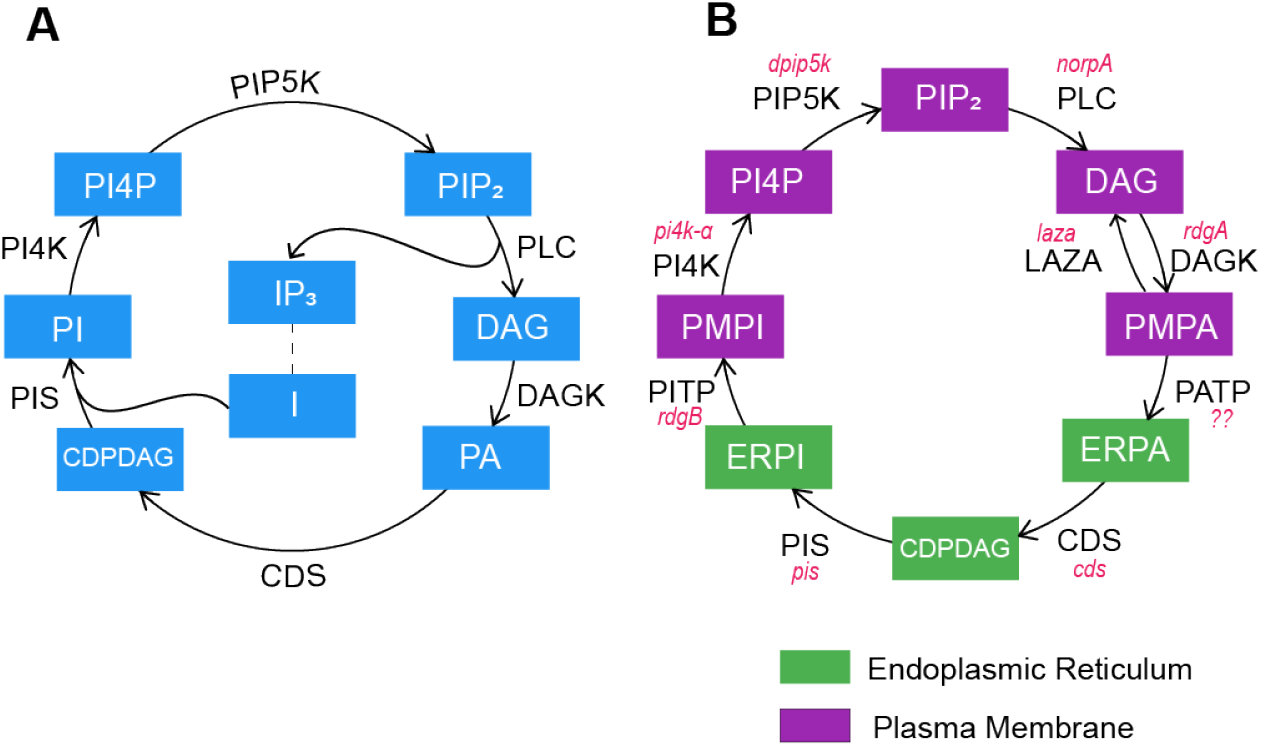
The PIP_2_ cycle. (A) Reactions for which there is direct experimental evidence of their involvement in phospholipase C (PLC) activated signalling in mammalian cells. Blue rectangles represent the chemical components in this cycle. Names in black represent the enzymes that catalyse the corresponding reactions. PLC hydrolyses phosphatidylinositol 4,5-bisphosphate (PIP_2_) into diacylglycerol (DAG) and inositol 1,4,5 trisphosphate (IP_3_). DAG is further hydrolysed by diacylglycerol kinase (DAGK) to form phosphatidic acid (PA) and then cytidine diphosphate-diacylglycerol synthase (CDS) synthesize cytidine diphosphate-diacylglycerol (CDPDAG). Phosphatidylinositol synthase (PIS) then condenses CDPDAG and myo-inositol to form phosphatidylinositol (PI). This PI is then used to resynthesis PIP_2_ by conversion of PI to phosphatidylinositol 4 phosphate (PI4P) by phosphatidylinositol 4-kinase (PI4K) followed by PI4P to PIP_2_ conversion via phosphatidylinositol 4 phosphate 5 kinases (PIP5K). (B) Similar cycle of reactions in Drosophila melanogaster photoreceptors. Here, components are distributed amongst the plasma membrane (PM; purple) and endoplasmic reticulum (ER; green). Known genes are labelled in italicized red text.

Many receptors trigger high and sometimes sustained levels of PLC activity. Examples of this include neuronal signalling mediated by metabotropic glutamate receptors or T-cell receptor signal transduction. Since PIP_2_ levels at the plasma membrane are low, cells need to rapidly restore its levels at this location during signal transduction so that PLC signalling is not hampered due to depletion of substrate. In doing this, cells face at least two challenges: (i) Co-ordinating the activities of each enzyme that is part of this multistep metabolic conversion; (ii) The lipid intermediates that are part of the metabolism of DAG are non-diffusible, yet the enzymes that participate in these reactions are located in distinct sub-cellular compartments (reviewed in [7]) (Fig 1 (B)). Prominent examples of this include the generation of DAG at the plasma membrane by PLC whereas CDS and PIS that are part of its metabolic conversion back to PIP_2_ are located on the endoplasmic reticulum. Further, the phosphoinositide kinases that phosphorylate PI to generate PIP_2_ are located at the plasma membrane. Thus, in addition, PIP_2_ resynthesis following PLC activation requires the co-ordinated transfer of lipid intermediates between these compartments. These considerations become especially important if the total amount of lipid intermediates present in a cell that can undergo PLC dependent PIP_2_ turnover is fixed; i. e. there is no injection of additional molecules of any intermediate into the cycle. We refer to this scenario as the “closed” PIP_2_ cycle. The biological relevance of such co-ordinated synthesis has been best articulated by the Li^3+^ hypothesis proposed by Berridge [4,5]. This hypothesis proposes that since Li^3+^ is an inhibitor of inositol 1 monophosphatase (IMPase), an essential enzyme involved in the recycling of inositol back to PI resynthesis, during Li^3+^ treatment of cells the resynthesis of PI, and consequently PIP_2_, is inhibited consequently downregulating PLC signalling. Crucially, the Li^3+^ hypothesis is underpinned by the assumption that the total amount of metabolic intermediates in the PIP_2_ cycle is fixed, although there is no experimental evidence to clearly establish this idea. Thus, an analysis of the nature of the PIP_2_ cycle and the co-ordination of turnover of intermediates is of crucial importance.

Signal transduction in *Drosophila melanogaster* photoreceptors are a typical example of G-protein coupled PLC signalling [17]. Photon absorption by rhodopsin triggers PLC activation leading to the hydrolysis of PIP_2_. The DAG and IP_3_ so produced then undergo metabolic conversion to regenerate PIP_2_ (Fig 1 (B)). The biochemical steps involved in the photoreceptor PIP_2_ cycle are orthologous to those described in a range of eukaryotic model systems including mammalian cells [7]. Drosophila mutants that affect most of the enzymatic steps in this cascade have been described. Importantly, biochemical analysis using both radiolabelling as well as mass spectrometry have reported changes in the levels of some of the key metabolic intermediates of the PIP_2_ cycle, especially DAG and phosphatidic acid (PA) [9,16,20]. Interestingly, it has previously been proposed that in Drosophila photoreceptors, the PIP_2_ cycle might not be closed but may involve the activity of a lipase that acts as a sink removing DAG from the cycle [6,13] although this remains unresolved (reviewed in [16,17]). Thus, PLC signalling in Drosophila photoreceptors offers a valuable model in the context of which to analyse the regulation of PLC activated PIP_2_ signalling, including a critical consideration of whether or not the PIP_2_ cycle is an “open” or “closed” cycle. In this setting, we have generated several mathematical models of the PIP_2_ cycle (See Materials and Methods) with a view to understanding whether it represents a closed or open cycle and the biochemical implications of either scenario. We find that current experimental data are not consistent with a mathematical model of a closed PIP_2_ cycle; rather these are explained by the existence of a sink coupled with a source that funnel metabolic intermediates through the PIP_2_ cycle. These results have fundamental implications for our understanding of PLC signalling, a process that underpins numerous basic cellular processes of biochemical importance.

## Results

### A closed PIP_2_ cycle is consistent with the wild-type steady-state data

The simplest scenario (as mentioned in Materials and Methods) is one where the PIP_2_ cycle is ‘closed’, i.e., no external sources or sinks exist that add or remove any of the intermediate lipids components of the cycle, and there are only ‘forward’, irreversible, first order reactions. In this scenario, the equations governing the dynamics of the lipid concentrations are linear (see equations S1 Eq-S8 Eq in supplementary section S1) and are fully specified by 8 reaction rate constants. We find that (see supplementary section S1) there are in fact an infinite set of possible of parameter (i.e. rate constants) values each of which are consistent with the experimentally observed data shown in Table 2. For each one of these infinite parameter sets, in the steady state in wild-type cells approximately 3.92% of the total lipid would consist of PIP_2_, 3.92% of PI4P, 0.63% of DAG, 13.14% of PA (counting both ER and PM), 78.33% of PI (counting both ER and PM), and 0.08% of CDP-DAG (see supplementary section S1). Thus, in wild type photoreceptors, a closed PIP_2_ cycle with only forward, irreversible, first-order reactions is consistent with the observed steady state levels of the lipid intermediates of the PIP_2_ cycle. We also find a similar result from *in silico* analysis of the Michaelis-Menten model of this closed PIP_2_ cycle (see supplementary section S6).

### Deviations from experimental data in the models of a closed PIP_2_ cycle

However, we find that the first-order and Michaelis-Menten models of the closed PIP_2_ cycle are inconsistent with experimental data from two mutants (supplementary section S1 and S6). First, in *rdgA*^3^ mutants it has been observed that the ratio of DAG concentration to total PI concentration is no different from that in the wild-type [11]. In contrast, our models of a closed PIP_2_ cycle predict a clear increase in this ratio when DAGK activity is decreased. In the model with first-order reactions, the increase is proportional to the decrease in DAGK activity in the mutant (see supplementary section S1; for a similar result with Michealis-Menten type kinetics, see supplementary section). Second, in mutants lacking the PA phosphatase LAZA *(laza*^22^), the ratio of total PA concentration to total PI concentration increases by > 200%, compared to the wild-type control [9]. We repeated experiments to measure [PA]_total_ levels in *laza^22^* and were able to reproduce the elevations in this ratio reported in earlier studies [20]. However, in our models, strong hypomorphs of LAZA exhibit no significant change in this ratio (see Table 1 and supplementary section S1). This inconsistency between the model prediction and experimental data implies that one or more assumptions in the model are not valid and the PIP_2_ cycle cannot be the simple closed cycle depicted in Fig 2(A).

**Figure 2.**
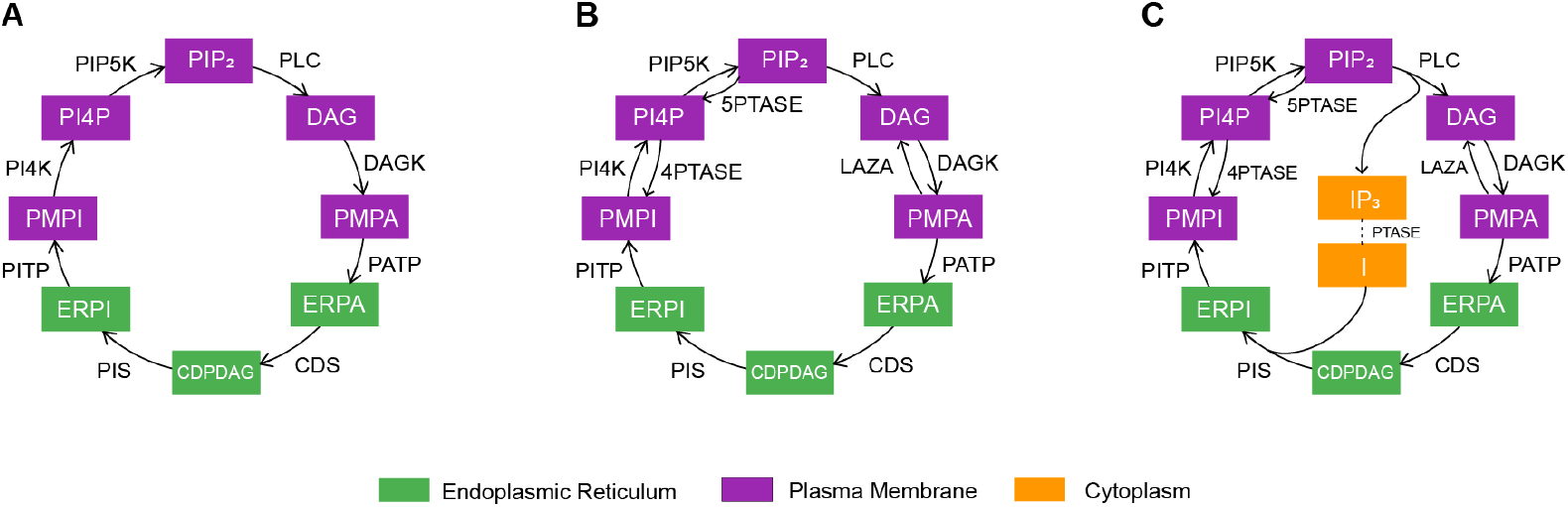
Closed PIP_2_ cycles. , where the total amount of lipid must remain constant. None of these are consistent with experimental observations.(A) Simplest case, containing only the ‘forward’ reactions. (B) Cycle with three additional ‘reverse’ reactions. For one of these, mediated by LAZA, there is direct experimental evidence in *Drosophila melanogaster*, while for the other two there are potential candidates. (C) ‘Classical’ PIP_2_ cycle with reverse reactions and explicit conversion of IP_3_ to myo-inositol, as well as the incorporation of myo-inositol in the PI synthesis reaction. There is no direct evidence for these reactions in*Drosophila melanogaster*, but they are known to exist in the mammalian PIP_2_ cycle.

**Table 1.** Comparison of experimental observations for various mutants with theoretical predictions from our models. Mismatches between experiments and theoretical predictions are indicated in red. The Open Cycle 2 configuration of Fig.(B) is the only one that is consistent with all experimental observations. All numbers are relative to the value of that ratio in wild-type cells, and are therefore dimensionless. >> 1 indicates that all parameter values that are consistent with the wild type steady state condition result in a value for that ratio that is much larger than unity. 1 indicates that all parameter values that are consistent with the wild type steady state condition result in that ratio being exactly unity. ≈ indicates that there exists at least one parameter set that is consistent with the wild type steady state condition and results in a value of that ratio that is approximately the value mentioned (see supplementary figure S7 Fig for the results from a range of parameter values).

|  | *rdgA*<sup>3</sup> |  | *laza*<sup>22</sup> |  |
| --- | --- | --- | --- | --- |
|  | *DAG/PI*<sub>total</sub> | *PA*<sub>total</sub>/*PI*<sub>total</sub> | *DAG/PI*<sub>total</sub> | *PA*<sub>total</sub>/*PI*<sub>total</sub> |
| Experimental observation (Reference) | **1** [11] | **1** [16], **0.98** [9] | **1** [9] | **2.49** [9] |
| Closed irreversible cycle | >> 1 | 1 | N/A | N/A |
| Closed reversible cycle | >> 1 | 1 | ≈ 1 | 1 |
| Closed classical cycle | >> 1 | 1 | ≈ 1 | 1 |
| Open cycle 1 | >> 1 | 1 | ≈ 1 | ≈ 2.49 |
| Open cycle 2 | ≈ 1 | 1 | ≈ 1 | ≈ 2.49 |

### Known reverse reactions are insufficient to recapitulate elevated PA/PI ratio in **laza^22^**

In addition to the eight forward reactions, there are in principle three known lipid phosphatases that can act on lipid components of the PIP_2_ cycle. These include a 5-phosphatase (that converts PIP_2_ to PI4P) [1,14], a 4-phosphatase (that converts PI4P to PI) [10] and a PA phosphatase that is able to dephosphorylate PA to generate DAG [9]. These three enzymes are able to render three of the forward reactions in the PIP_2_ cycle reversible. We found that adding these three reactions to our models of the closed PIP_2_ cycle did not alter our results - the models were consistent with wild-type steady state observations, but not with observations reported for specific mutants (supplementary section S2). In particular, these models are also unable to explain the increase in PA/PI ratio experimentally observed in *laza*^22^.

### Recycling of myo-inositol is also insufficient to recapitulate elevated PA/PI ratio in **laza^22^**

The third model we investigated adds one more known element to the intermediates of the PIP_2_ cycle: In addition to DAG, the activity of PLC on PIP_2_ also produces IP_3_ as a second product. IP_3_ is converted to myo-inositol in a three-step process following which the enzyme PIS condenses myo-inositol to CDP-DAG to form PI on the endoplasmic reticulum. Thus, in a situation where the PLC catalysed production of DAG is very high, it is possible for PI production from CDPDAG to be substrate limited due to the absence of sufficient myo-inositol. This is a potential effect that our previous models ignored. However, including these steps too cannot explain (see supplementary section S3) the large increase of the PA/PI ratio observed experimentally in the *laza^22^* mutant.

### Feedback loops cannot recapitulate elevated PA/PI ratio in *laza^22^*

There is experimental evidence to support the existence of feedback loops within the cycle, where a lipid component may affect the activity of an enzyme that is part of the cycle. For example, the activity of PIP5K has been reported to be enhanced with increasing PA levels [15]. Such feedback can be modelled by making the enzyme activity a function of the levels of this lipid component. For example, in the model where the reactions are first-order, the effect of PA on PIP5K could be represented as follows:

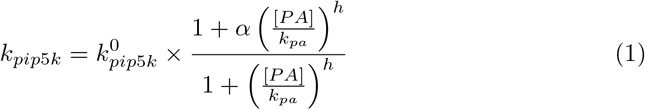

where 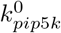 is the PIP5K activity in the absence of PA, *α* is fold-increase in the PIP5K activity in the presence of saturating amounts of PA, *h* is the Hill coefficient (a measure of the cooperativity of the feedback), and *k_pa_* sets the concentration of PA at which the feedback kicks in. For these feedback models, every parameter set that produces a valid steady-state for the cycle without feedback has a corresponding parameter set that produces exactly the same steady-state for the cycle with the feedback link (see supplementary section S3). Therefore, a closed cycle with feedback links will not explain the steady-state [PA]_total_ /[PI]_total_ ratio in *laza^22^* any better than the cycle without feedback.

### An open cycle topology can recapitulate PA/PI increase in *laza^22^*

Our results thus far suggest that the PIP_2_ cycle is not a closed cycle, i.e., additional lipid must be entering the cycle from external biochemical reactions not included in the closed cycle models, which we will term “sources”. In order to achieve a steady-state, there must then also be one or more processes (“sinks”) that balance the external sources by removing lipid from the cycle. We explored all possible configurations of the PIP_2_ cycle which had one source and one sink somewhere in the cycle and asked which source-sink combination was able to recapitulate the elevated PA/PI ratio observed in *laza^22^*. We found that indeed some configurations of such an open cycle were able to recapitulate the increased PA_total_/PI_total_ ratio in **laza^22^**, while simultaneously being consistent with the observed wild-type lipid ratios of Table 2. In these successful configurations, the sink could be in only one of two locations, as shown in Fig 3 (A) and (B). The first possibility for the sink is the PA phosphatase activity of LAZA acting at the plasma membrane and generating a pool of DAG that is no longer able to re-enter the PIP_2_ cycle (Fig 3 (A)). The second possibility is an enzyme, or DAG binding protein, that metabolizes DAG into a product that cannot be recovered back into the PIP_2_ cycle (Fig 3 (B)). With either one of these sinks operational, it is possible to recapitulate in our model the elevation of PA_total_/PI_total_ ratio seen in **laza^22^**. However, the former sink, acting on PA at the plasma membrane, is inconsistent with the experimental observation that the DAG/PI_total_ ratio remains constant in *rdgA^3^* mutants that lack DAGK activity (see supplementary section S4; this result holds irrespective of the location of the source). With the latter sink, namely that based on an enzyme or binding protein capable of metabolizing DAG, it is possible to recapitulate both an elevated PA/PI ratio in *laza*^22^ as well as the constancy of the DAG/PI ratio in *rdgA*^3^ (more precisely, with first-order reactions we found parameter sets that exhibited about 2-fold increase in PA/PI, see supplementary section S5, while with Michaelis-Menten reactions we found parameters that exhibited 2.5-fold and higher increases in this ratio in **laza^22^**, see supplementary section S6). Thus, this open cycle with Michaelis-Menten reactions is in fact consistent with all the wild-type and mutant steady-state experimental observations we have used in this paper (Table 1 summarizes the status of all the cycles we have studied). Note that, in this case, the location of the source at ERPA is necessary; configurations with the source at other locations are not consistent with the experimental observations (see supplementary section S5).

**Figure 3.**
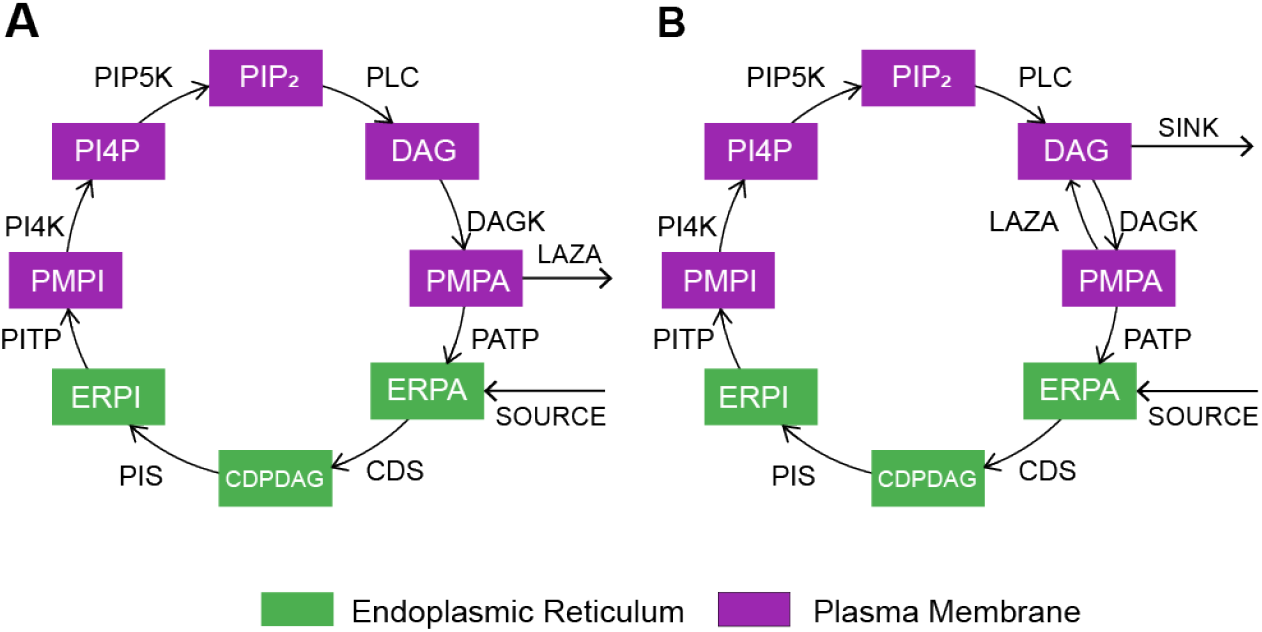
Open PIP_2_ cycles containing one sink and one source, that are capable of exhibiting the> 200% rise observed in PA/PI in *laza^22^* mutant. (A) Open cycle 1, where LAZA converts PA on the plasma membrane to a pool of DAG which is unable to further participate in the PIP_2_ cycle. This configuration is unable to explain the absence of increase in DAG/PI ratio in an *rdgA^3^* mutant (this result holds for all locations of the source). (B) Open cycle 2, where a hypothetical enzyme on the plasma membrane metabolizes or degrades DAG thus removing it from the PIP_2_ cycle. This configuration is able to explain the constancy of DAG/PI ratio in the *rdgA*^3^ mutant. Here, the location of the source at ERPA is crucial; other locations are incompatible with the experimental observations (see supplementary section S5).

**Table 2.** Ratios of lipid intermediate concentrations used in our mathematical models for the steady state condition in wild type.

| Lipid Ratio | Value | Reference |
| --- | --- | --- |
| $[PIP_2]/[PI]_{total}$ | $\approx 0.05$ | [2, 12, 19] |
| $[PI4P]/[PI]_{total}$ | $\approx 0.05$ | [2, 12, 19] |
| $[DAG]/[PI]_{total}$ | $\approx 0.008$ | Measurement from R. Padinjat's lab |
| $[PA]_{total}/[PI]_{total}$ | $\approx 0.1677$ | Measurement from R. Padinjat's lab |
| $[CDPDAG]/[PI]_{total}$ | $\approx 0.001$ | [21] |

## Discussion

The robustness of any signalling system depends on the availability of adequate substrate to fuel the rate limiting biochemical reactions that form part of this pathway. Depletion of substrate would be detrimental leading to slowing down of rate limiting steps and degrading the information context that is being conveyed. The PLC mediated signalling system that is used in diverse biological contexts in eukaryotes is triggered by PLC activation by cell surface receptor occupancy and in this context the adequate availability of its low abundance substrate PIP_2_ is a key requirement for sustained signalling. This process is particularly important in the context of cellular process such as in neurons where neurotransmitter receptors (such as the metabotropic glutamate receptor) trigger high rates of PLC activation. The observation that PIP_2_ levels undergo only transient, if any, depletion during cell signalling [8] implies that robust biochemical processes underpin the ability of cells to support its levels at the plasma membrane.

What might these processes be? The synthesis of PIP_2_ in eukaryotic cells is mediated by the sequential phosphorylation of hydroxyl groups at positions 4 and 5 of the inositol ring of PI. Thus, the availability of adequate amounts of PI is a minimal requirement for the resynthesis of PIP_2_ that has been depleted by PLC activity. Measurements by mass assays reveal that the mass of PI is about 20 times the levels of either PI4P or PIP_2_. While this might imply that there is a vast excess of PI from which to generate PIP_2_, it is important to note that most of this PI is located in the ER a distinct compartment from which it must be made available at the plasma membrane where the kinases that generate PIP_2_ are located. The mass of PI at the plasma membrane is unknown and has remained difficult to determine experimentally although it is widely accepted that the vast majority of the PI in a eukaryotic cell is at the endoplasmic reticulum. Thus, it is very likely that the pool of PI available at the PM for PIP_2_ resynthesis is relatively limited.

PI is synthesized by the condensation of CDP-DAG with Inositol by the enzyme phosphatidylinositol synthase. The availability of substrates for this reaction may arise from one of two sources: (i) they may be generated as downstream metabolites generated by the primary products of PLC activity, i.e. IP_3_ and DAG. IP_3_ is sequentially dephosphorylated to generate Inositol, and DAG is sequentially converted into PA and then CDP-DAG. Thus, in the context of a closed cycle, i.e. one with no exogenous source of PI, the control of DAG and PA levels during PLC signalling is vital to support PIP_2_ resynthesis. (ii) PI may be generated by use of de novo synthesized CDP-DAG and inositol that is transported into cells from the extracellular medium. In this setting, which is an example of an open cycle, the conservation and recycling of metabolic intermediates following PLC activity is likely to be less important to maintain PIP_2_ levels through resynthesis.

To date there have been no biochemical measurements/studies that have conclusively defined the PIP_2_ cycle during PLC activation as either an open or a closed cycle. Yet the closed/open nature of this cycle has important functional implications for biological processes underpinned by this PLC signalling system. As an example, the Li^3+^ hypothesis of Berridge, which proposes that the inhibition of IMPase, a key enzyme in the cycling of IP_3_ to inositol during PLC signalling, leads to the depletion of PI and consequently PIP_2_, implicitly assumes that there is no additional source of inositol for PI synthesis of PI during cell signalling. Likewise, a recent study proposed that the levels of PA regulated by the activity of DAGK and PA phosphatase might limit the resynthesis of PIP_2_ in Drosophila photoreceptors during PLC signalling [9]. An open cycle with external sources necessitates a re-evaluation of these hypotheses; an additional source of inositol could decouple the rate of PI synthesis from the activity of IMPase, and an external source for PA could similarly counteract changes in the activity of DAGK and PA phosphatase.

In this study, we have constructed a quantitative mathematical model of the PIP_2_ cycle. At steady state, as evaluated by the ratios of DAG/PI and PA/PI, this model recapitulates experimentally measured values of the ratios for these key lipids in the PIP_2_ cycle. Although an infinite parameter space was able to recapitulate these ratios for wild type cells, we found that when interrogated for the impact of specific mutants, the model revealed anomalies with respect to the levels of PA and DAG experimentally reported to arise from the manipulation of DAGK and PA phosphatase activity. These anomalies could not be resolved at any point in the parameter space assuming a closed cycle. However, the experimentally noted elevation of PA levels arising from the loss of PA phosphatase could be recapitulated *in silico* by assuming the existence of an additional source of PA as well as a sink. It was necessary to assume a sink at one of two locations: (i) A PA phosphatase activity at the plasma membrane that generates DAG; this pool of DAG being no longer able to participate in the PIP_2_ cycle (Fig 3 (A)), (ii) a DAG metabolizing activity at the plasma membrane that is able to convert DAG into metabolites that can no longer participate in the PIP_2_ cycle (Fig 3 (B)). While both of these sinks, in conjunction with a source can be invoked to explain the accumulation of PA in the PA phosphatase mutant, the presence of a PA phosphatase activity that creates an unavailable pool of DAG should have resulted in an experimentally measurable elevation in the levels of DAG; such elevations have not been demonstrated so far. By contrast, the presence of a DAG metabolizing activity at the plasma membrane, would provide a parsimonious explanation for lack of elevation of DAG levels when DAGK activity is reduced. Although candidate DAG lipases that function in Drosophila photoreceptors in the context of PLC signalling have been proposed, the enzyme has not been definitively identified. If such a mutant in DAG lipase were isolated, our model predicts that DAG levels would likely be elevated during PLC signalling. In mammalian models, loss of DGK-є has been reported to result in shorter tonic seizures and is associated with accumulation of arachidonic acid that can be generated from DAG by DAG lipase [18].

In our mathematical model, the presence of a sink must be matched by a source; otherwise the total mass of metabolites in the cycle will reduce rapidly. While we have not explicitly examined the case of multiple sources (or indeed multiple sinks), our model predicts that there must at least be one sink at DAG on the plasma membrane and one source that injects PA at the endoplasmic reticulum. Our model cannot say whether this is due to de novo synthesis of PA or its generation by the activation of phospholipase D (PLD). Numerous experiments have shown that cell surface receptors that activate PLC often also activate PLD. The significance of PA generated by PLD as a potential source for the PIP_2_ cycle needs to be evaluated. Nonetheless, our results highlight the need to establish the sources and sinks that may provide additional points of control for this key signalling cycle.

## Materials and Methods

### Construction of mathematical models of the PIP_2_ cycle

We made several variants of a mathematical model of the PIP_2_ cycle, based on the well-known reaction cascade shown in Fig 1 (B). The models consist of differential equations describing the rate of change in concentration of each individual component of the cycle. Importantly, given that the lipid intermediates of the cycle cannot diffuse across the cytosol and are therefore restricted to the membrane at which they are produced, we model the lipid intermediates that are found both at the plasma membrane and the endoplasmic reticulum as separate components with separate equations. In addition, we include reaction rates for the specific transfer of lipid intermediates from one compartment to the other by lipid transfer reactions.

The models constructed differ in the way we mathematically represent the reaction rates, as indicated in Fig 4. The simplest model assumes that all reactions are irreversible and of first-order, and there are no external sources or sinks, as shown in Fig 2(A); thus, for example, the rate of change of PIP_2_ concentration is described by the following equation,

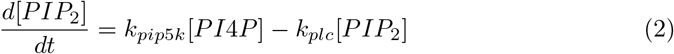

where the first term on the right-hand side represents the rate of synthesis of PIP_2_ from PI4P, and the second term represents conversion of PIP_2_ to DAG, with *k_pip5k_* and *k_plc_* being the corresponding first-order rate constants (the equations for other components are similarly formed by summing all synthesis and consumption reactions for a given substrate - see supplementary section 2 for the full set of equations). The next variant of the model also uses only first-order reactions and no external sources or sinks, but adds three ‘reverse’ reactions, as shown in Fig 2(B). Further variants use the reaction schemes of Fig 2(A) and Fig 2(B), but replace the first-order reactions by Michaelis-Menten reactions. Thus, instead of a single reaction constant describing each synthesis and consumption reaction, these variants have two parameters, the maximum reaction rate and the Michaelis constant, as shown in Fig 4. Finally, we constructed models that allowed one external source and one sink which, respectively, add or remove lipid from the cycle, as shown in Fig 3(A) and (B). These models were examined both with first-order, as well as with Michaelis-Menten reactions.

**Figure 4.**
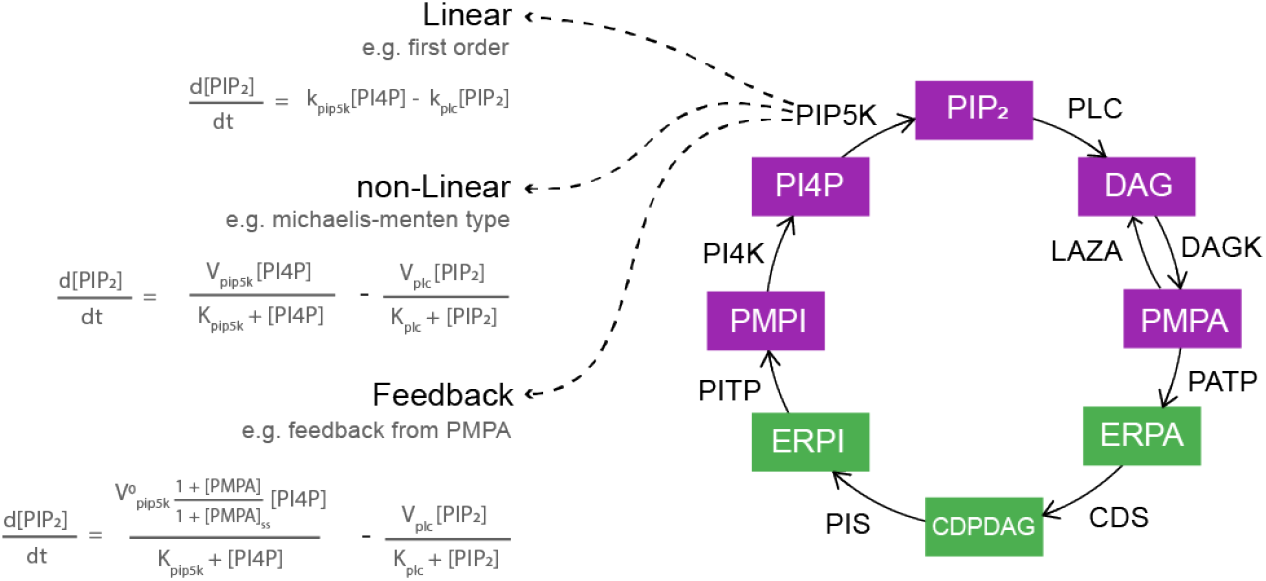
Representative mathematical forms used to model the kinetics of individual chemical components of the PIP_2_ cycle. See supplementary sections S1-S6 for complete lists of reaction rate equations.

### Steady-state analysis of the models

At steady state, the rate of change of concentration of each component in the cycle will be zero. For example, equation 2, above, implies that the steady state concentration of PI4P will be related to the steady state concentration of PIP_2_ as follows:

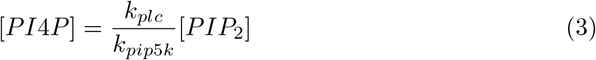

Similarly, we can show, for this simple first-order reaction model, that in steady state the ratio of PIP_2_ to PI_total_ lipid concentration is given by (see supplementary section):

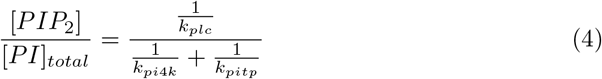

Thus, changes in the activities of any of the three enzymes, PLC, PI4K or PITP will affect the steady state ratio of PIP_2_ to PI_total_, but changes in PITP will not affect the ratio of PIP_2_ to PI4P. Such formulae, along with experimentally determined information about the ratios between individual lipids at steady state (see Table 2), allow us to put strong constraints on allowed values for the reaction rate constants in the above equations. In particular, if we cannot find any values for the set of reaction rate constants that satisfy the experimentally observed lipid ratios in Table 2, then we can infer that that model is incompatible with the observed data. For models using the first-order reactions, this comparison can be done analytically (supplementary sections S1-S5). For models using Michaelis-Menten reactions, we perform this analysis numerically, *in silico* (supplementary section S6).

## Acknowledgments

We thank members of the Padinjat lab for sharing unpublished data and measurements and for numerous discussions. This work was supported by the National Centre for Biological Sciences-TIFR, the Simons Foundation and a Wellcome-DBT India Alliance Senior Fellowship to RP.

## Supporting information

**S1. Closed PIP_2_ Cycle with only Irreversible, First-Order Reactions**

**S2 Closed First-Order PIP_2_ Cycle with Three Additional Reverse Reactions**

**S3 Classical Closed First-Order PIP_2_ Cycle**

**S4 Open Cycle 1** Sink which removes PMPA from the cycle, catalysed by LAZA

**S5 Open Cycle 2** Sink which removes DAG from the cycle

**S6 Michaelis-Menten Kinetics**

**S7 Fig. Fold-changes in DAG/PI**_*total*_ **(red) and PA**_*total*_/**PI**_*total*_ **(blue) ratios.** for the parameter sets we found that satisfied the wild type steady state constraints with an error of less than 15% (10559 for panel A, 3413 for panel B, 6311 for panel C and 14276 for panel D; a few of these points fall outside the axis ranges.). The x-axes show this fold-change in the *rdgA^3^* mutants, while the y-axes show the fold-change in the *laza*^22^ mutant (note that for the closed, irreversible cycle of Fig S1, we can only test the *rdgA^3^* mutant). The black arrows in (D) point to the parameter set that came closest to simultaneously satisfying all experimental observations; the corresponding parameter value, and the resultant fold-changes in lipid ratios are shown in Fig S8.

**S1 Eq. Rate of change of PIP_2_ concentration**

**S8 Eq. Rate of change of PI4P concentration**

